## Supplemental Material for "Study of changes in brain dynamics during sleep cycles in dogs under effect of trazodone"

June 16, 2025

#### S1. Correlation between REM latencies with Age and Weight

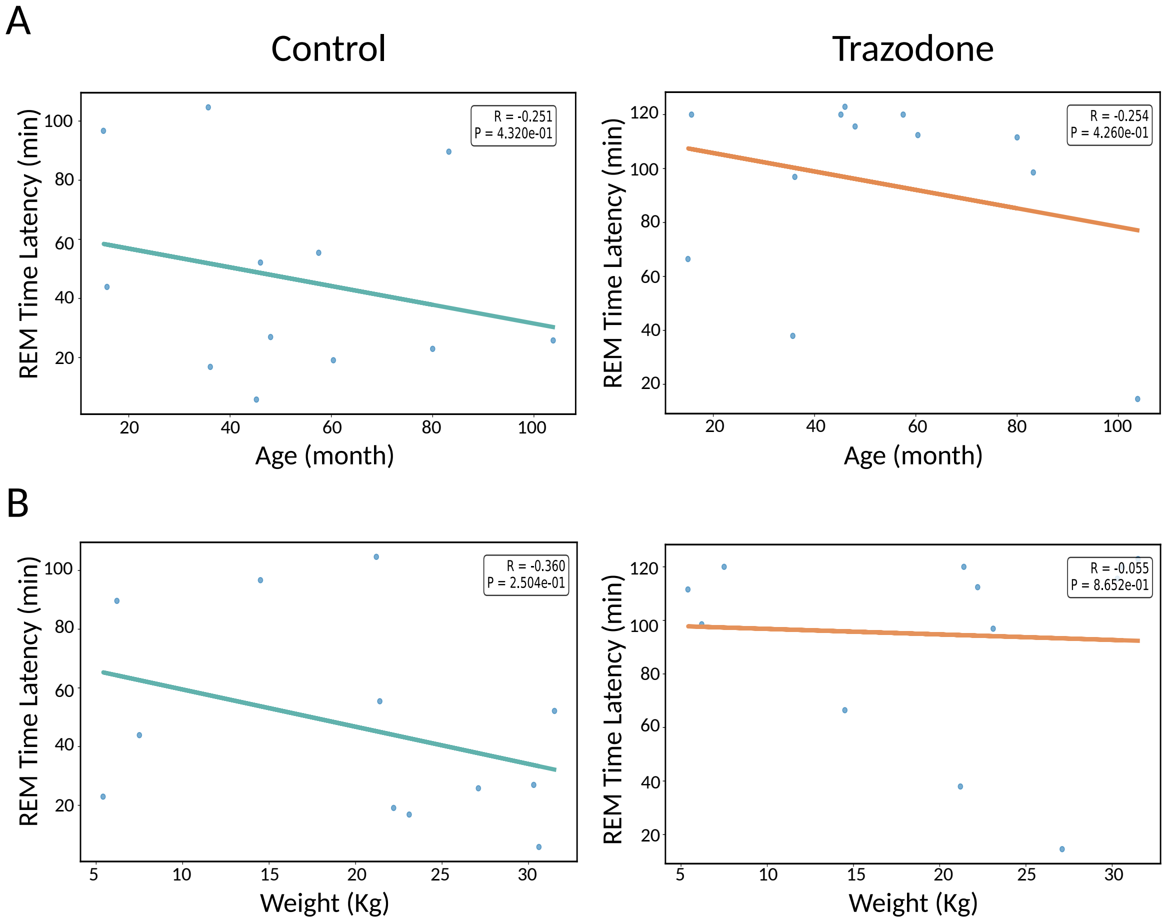


Figure 1: Correlation between REM latency with A) Age and B) Weight.

### S2. Sleep transition Matrix

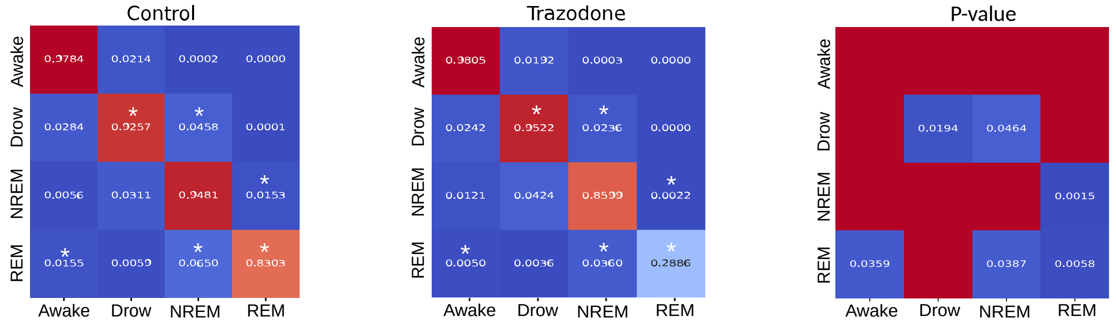


Figure 2: Transition matrices between different sleep states for Control and Trazodone conditions (left and center, respectively). Asterisks indicate statistically significant differences. $p$-values represent comparisons between matrices in both conditions. Statistical analysis was performed using the Kruskal–Wallis test followed by Dunn's post hoc correction.

### S3. Coherence Analysis

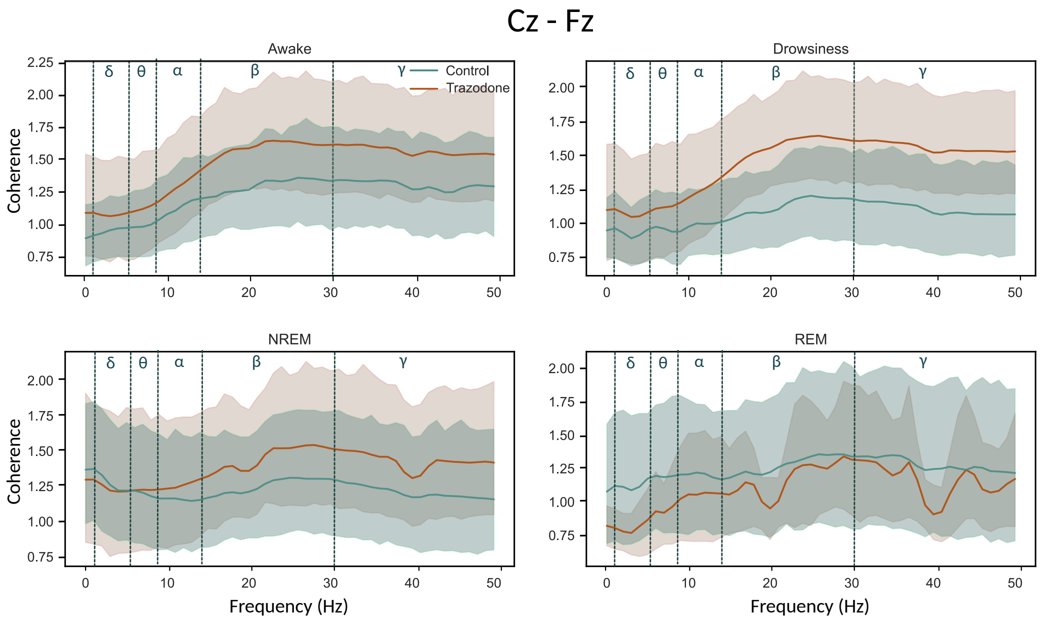


Figure 3.1. Coherence analysis for Cz-Fz electrode pairs across all behavioral states. The green and red traces denote the control and trazodone groups, respectively. Vertical blue dashed lines indicate boundaries of standard physiological frequency bands (delta $\delta$, tetha $\theta$, alpha $\alpha$, beta $\beta$, gamma $\gamma$). Observed attenuations near 20 Hz and 40 Hz likely reflect harmonic suppression by the notch filter.

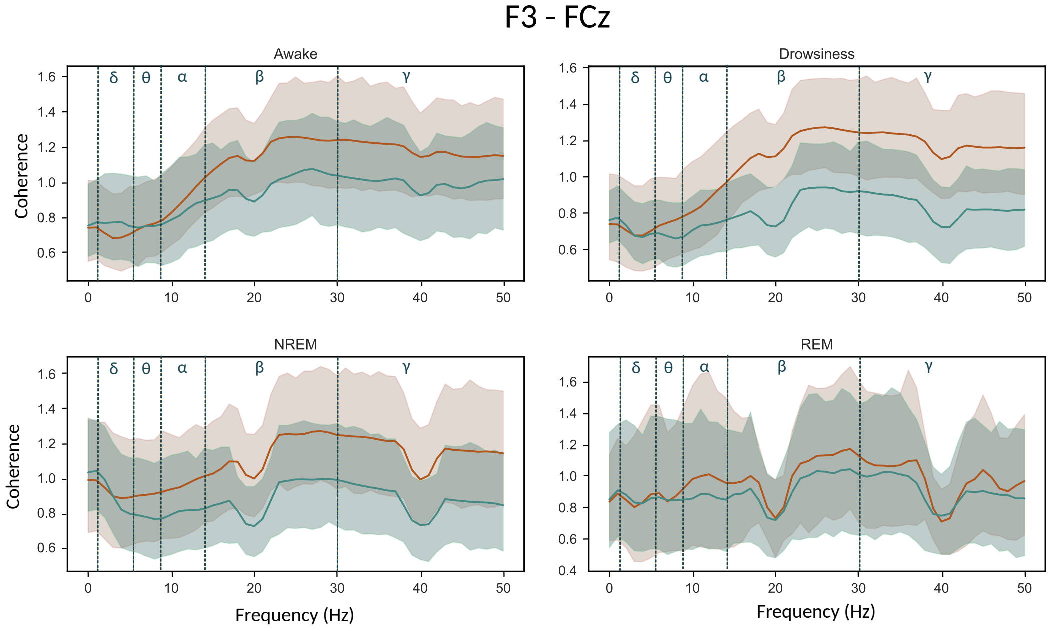


Figure 3.2. Similarly to Figure 3.1 for F3-FCz electrode pairs.

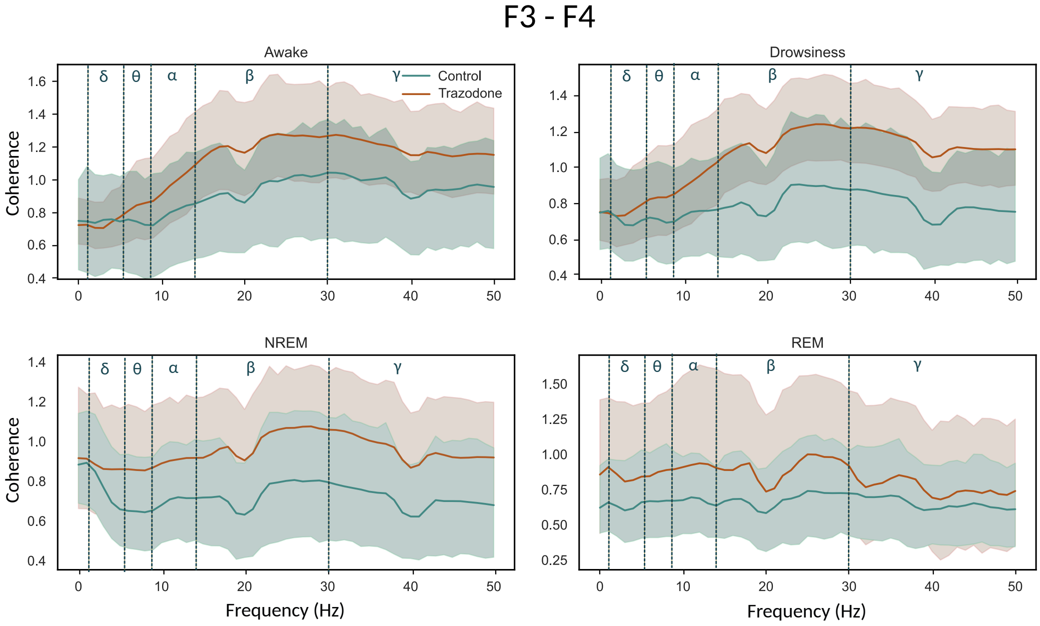


Figure 3.3. Similarly to Figure 3.1, for F3-F4 electrode pairs

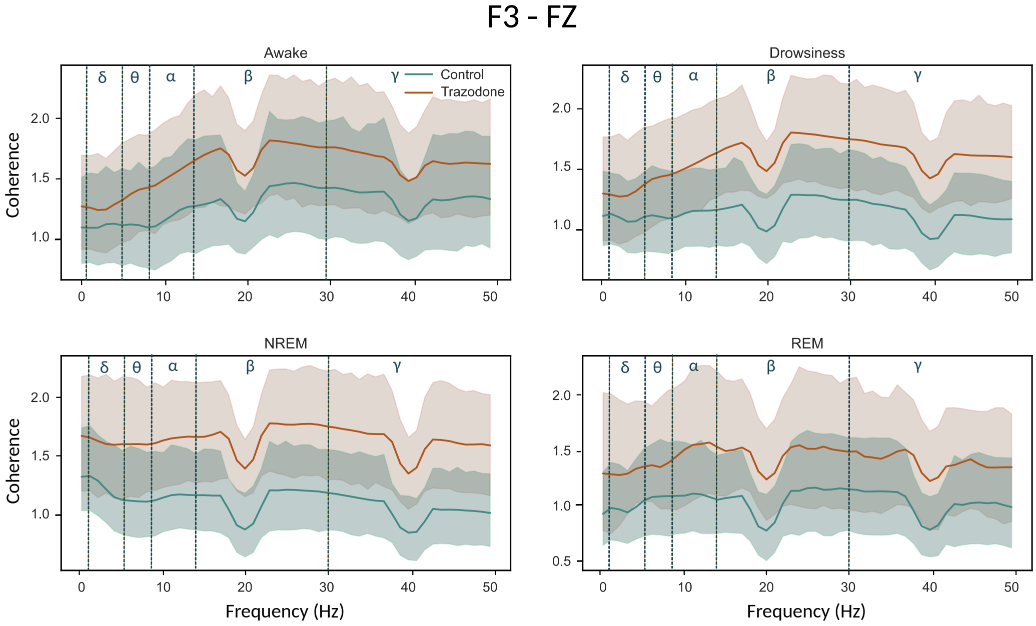


Figure 3.4. Similarly to Figure 3.1, for F3-Fz electrode pairs

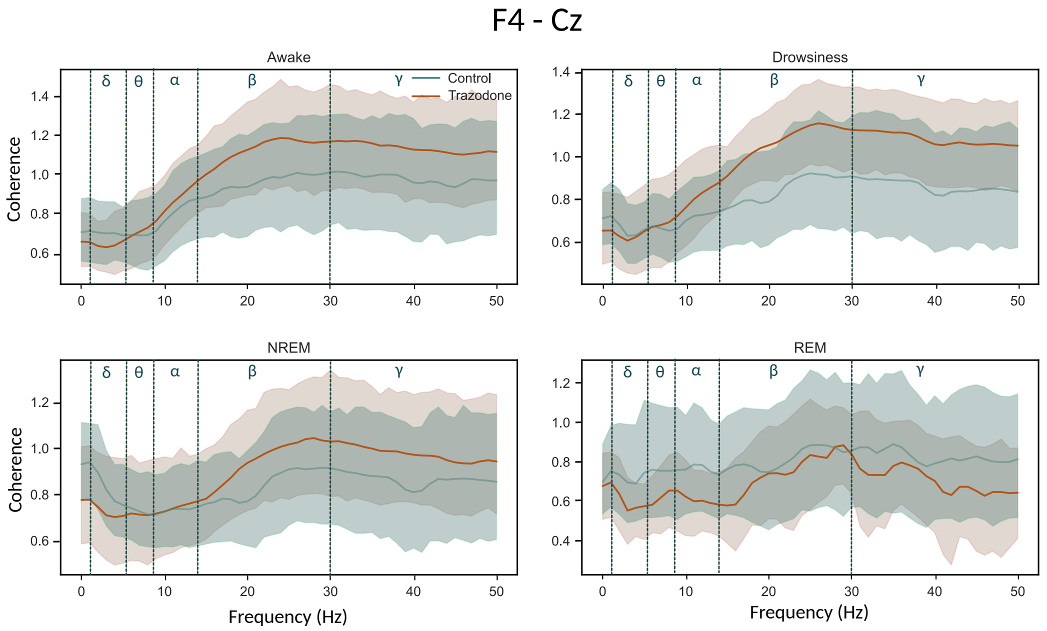


Figure 3.5. Similarly to Figure 3.1, but for F4-Cz electrode pairs

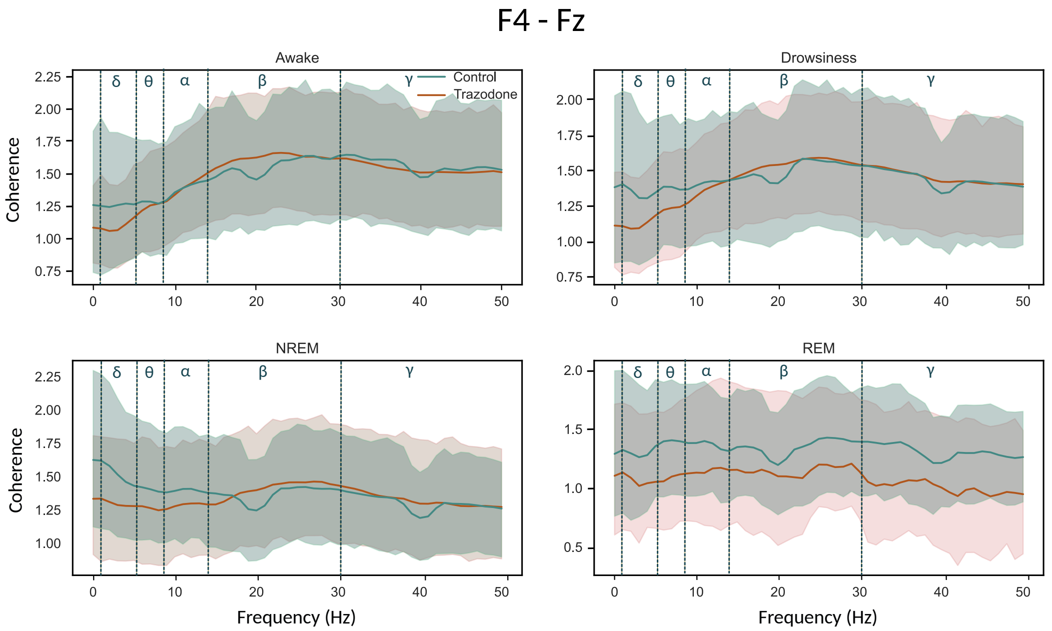


Figure 3.6. Similarly to Figure 3.1, but F4-Fz electrode pairs
